# Chlamylipo, a *Chlamydomonas*-in-liposome microswimmer: self-propelled swimming and associated lipid membrane flow

**DOI:** 10.64898/2026.03.11.711009

**Authors:** Shunsuke Shiomi, Koichiro Akiyama, Hiromasa Shiraiwa, Sota Hamaguchi, Daiki Matsunaga, Tomoyuki Kaneko, Masahito Hayashi

**Affiliations:** Department of Frontier Bioscience, Graduate School of Science & Engineering, Hosei University, Koganei, Tokyo 1848584, Japan; Graduate School of Engineering Science, The University of Osaka, Toyonaka, Osaka 560-8531, Japan; Department of Biotechnology and Life Science, Tokyo University of Agriculture and Technology, Koganei, Tokyo 184-8588, Japan

**Keywords:** *Chlamydomonas reinhardtii*, liposome, membrane flow, microswimmer, biohybrid robots

## Abstract

Developing active transport systems for microcargo delivery is challenging and requires overcoming the low Reynolds number constraints. We developed a bio-hybrid micro-swimmer, “chlamylipo” consisting of the green alga *Chlamydomonas reinhardtii*, encapsulated within a giant liposome. Although internal encapsulation offers cargo protection, it requires a mechanism to transmit the propulsion force across a closed membrane. We demonstrated that chlamylipo exhibited forward swimming and phototactic directional control. High-speed imaging of membrane shape and fluid flow revealed that the driving force originated from periodic membrane deformations and was accompanied by characteristic fluid dynamics. Flow analysis showed rapid oscillations at tens of hertz corresponding to flagellar beating, superimposed on slower axial migration at approximately 4 Hz associated with cell rotation. Corresponding flow signatures were also detected in the external fluid, indicating mechanical coupling across the lipid bilayer. Membrane domain tracking further showed that fluid motions inside and outside the membrane were coupled through viscous friction and membrane deformation, generating a characteristic four-vortex flow field consistent with a two-point force model. Together, these results suggest that membrane flow mainly reflects force transmission across the bilayer, whereas forward propulsion is primarily driven by periodic membrane deformation. This study elucidates the physical mechanism of force transmission in encapsulated swimmers, demonstrating that internal hydrodynamic power can effectively drive the motion of macroscopic containers.

**Significance:** The development of autonomous micro-swimmers for targeted drug delivery is a major challenge in biophysics. We present “chlamylipo,” a hybrid system in which a swimming alga is encapsulated inside a lipid vesicle. This study is significant because it demonstrates that an enclosed swimmer can propel a macroscopic container solely via hydrodynamic coupling across a closed membrane without direct external mechanical links. Furthermore, we achieved external directional control using phototaxis. This study provides physical insights into fluid-membrane interactions and proposes a novel strategy for designing light-guided active transport carriers.

## Introduction

In recent years, liposome-based drug delivery systems (DDS) have been extensively investigated. Liposomes are artificial lipid vesicles composed of lipid bilayers that can encapsulate substances of various properties and sizes [1]. In passive DDS that utilize blood flow within the body, knowledge has accumulated regarding methods for adjusting liposome size to avoid clogging or elimination in the bloodstream, as well as surface modification techniques to allow targeted accumulation and binding at specific sites [2,3]. In environments without blood flow, such as the body surface, ocular cavity, digestive system, respiratory tract, or abdominal cavity, liposome size and surface characteristics are not constrained in the same way; however, a driving force is required for their active transport to the target location. Current research focuses on utilizing microorganisms as propulsive agents for the active transport of micro-substances. There are two main approaches: externally driven, in which microorganisms have microparticulate cargo attached to their surfaces for transport [4], and internally driven, in which microorganisms are encapsulated with cargo inside liposomes. Externally driven types are limited in terms of cargo size and properties and have a low loading capacity, but benefit from the direct propulsion of microorganisms [5,6]. There have been reports of using microorganisms such as *E. coli* and *Chlamydomonas reinhardtii*, with studies showing that *the phototaxis of Chlamydomonas* can be used to control the direction of movement. The internally driven type has the advantage of transporting substantial quantities of cargo with various properties. Liposomes encapsulating *E. coli* move as cohesive units [7]. However, to control the movement direction of liposomes, it is necessary to effectively utilize the taxis of these microorganisms. Although *E. coli* exhibits chemotaxis, chemoattractants and repellents cannot be effectively utilized because they are blocked by the liposomal membrane. Therefore, utilizing *Chlamydomonas*, an organism exhibiting phototaxis, offers a promising strategy for regulating movement direction through light stimuli that can penetrate liposome membranes, thereby addressing the critical challenge of directional control in internally driven systems [8-12].

*Chlamydomonas* swim forward by moving its two flagella in a breaststroke-like motion and exhibits phototaxis by changing its swimming direction in response to light stimuli [8-10]. The flagellar beat consists of a power stroke, in which the two flagella open wide and move from the front to the back, and a recovery stroke, in which the flagella are pulled close to the cell body and moved from the back to the front. This time-reversal asymmetric cyclical motion produces forward swimming in low-Reynolds number environments. Because the flagella move at a slight angle relative to the cell’s anterior-posterior axis and the flagellar plane containing the flagellar roots, the cell body rotates in the direction of a left-handed screw as it moves forward. The cell body has a single eyespot, which allows it to repeatedly sense light intensity in all directions as it rotates. By adjusting the beating of its two flagella in synchrony with changes in the received light intensity, *Chlamydomonas* achieves phototaxis, altering its swimming direction toward or away from light sources.

In this study, we demonstrated that liposomes encapsulating *Chlamydomonas* (“chlamylipo”) exhibit forward swimming and phototaxis. We found that membrane deformation is the driving force behind chlamylipo movement and identified a correlation in oscillation frequencies, suggesting a coupling between fluid motion inside and outside the membrane. Furthermore, we demonstrated that the unique membrane flow patterns generated by flagellar beating could be reproduced using a two-point force model that simulated flagellar action.

## Materials and methods

### Chemicals

The phospholipids 1,2-dioleoyl-*sn*-glycero-3-phosphocholine (DOPC), 1,2-dipalmitoyl-*sn*-glycero-3-phosphocholine (DPPC), 1,2-diphytanoyl-*sn*-glycero-3-phosphocholine (DPhPC), and 1,2-distearoyl-*sn*-glycero-3-phosphoethanolamine-N-(lissamine rhodamine B sulfonyl) (ammonium salt) (RhPE) were purchased from Avanti Polar Lipids (USA). DOPE and ATTO 488-DOPE (488PE) were purchased from ATTO-TEC. DOPC was used because of widespread use in liposome preparation. A mixture of DPhPC, DPPC, and cholesterol was used to generate phase-separated membrane domains for flow visualization. MCT oil (Nisshin OilliO) was purchased from supermarkets. Phospholipids dissolved in chloroform (10 mM or 10 μM) were stored at −20°C. MCT oil was degassed at 1.2 kPa for 30 min, flushed with nitrogen, and stored in the dark at room temperature. All other chemicals were purchased from Wako, TCI, or Merck.

### Cultivation of *Chlamydomonas*

Wild type (*cc124* and *cc125*), paralyzed flagella mutant (*pf18*), and impaired flagellar autotomy mutant (*fa1*) of *Chlamydomonas reinhardtii* were gifted from Prof. Masafumi Hirono (Hosei University). *Chlamydomonas* was cultured in TAP medium at 26^°^C with aeration and 12 h/12 h light/dark cycles.

### Preparation of lipid solutions

Phospholipids dissolved in chloroform were mixed in a PCR tube, and the total lipid content in the tube was set to 100 nmol. The lipid composition was set to DOPC:Atto488PE = 1:0.01 (molar ratio). Chloroform was completely evaporated in a desiccator at 1.2 kPa for 30 min. After adding 100 μL of MCT oil to the dried lipids, the mixture was heated at 70°C whilst inverting and vigorously tapping the tube every 5 minutes to obtain a clear 1 mM solution. PCR tubes containing lipid solutions were stored in a dry atmosphere with silica gel in the dark at room temperature and used within six months. Before use, lipid precipitation was checked using phase contrast and fluorescence microscopy. To visualize membrane flow using membrane phase separation, the lipid composition was set to DPhPC:DPPC:Atto488PE = 0.5:0.5:0.01 (mol), and 100 μM water-soluble cholesterol was added to the external solution before observation, inducing the formation of phase separation domains in the membrane.

### Encapsulation of *Chlamydomonas* into giant unilamellar vesicles

*Chlamydomonas* cells were encapsulated into giant unilamellar vesicles by W/O emulsion-transfer method with centrifugation [13,14]. Briefly, cultured cells in the logarithmic growth phase were harvested and resuspended in an inner solution (100 mM sucrose, 30% Percoll, and 1 mg/ml BSA) at 1.5x10^7^ cell/ml. Next, 2 μl of the inner solution was added to 20 μl of the lipid solution in a PCR tube. The tube was agitated vigorously by sliding it on a tube stand several times to obtain a whitish emulsion. The entire emulsion was placed on 30 μl of outer solution (100 mM glucose, 1 mg/ml BSA) in another PCR tube. The tube was centrifuged at 500 G and 20 °C for 5 min to generate chlamylipos and precipitate them at the bottom of the tube. The oil layer was removed, and the precipitate was collected with the outer solution and resuspended in a new tube by tapping.

### Microscope imaging

The motion of chlamylipos was observed using phase contrast, dark field, and epifluorescent microscopes (BX53 and IX71, Olympus, Japan) equipped with a 40x objective lens. Simultaneous observation of the autofluorescence of the cell body (red) and the fluorescence of the liposome membrane (green) was achieved using a wide-pass dichroic mirror unit (U-FBW, Olympus, Japan). The images were captured using a high-speed CMOS camera (DFK33UX287 for color and DMK33UX287 for monochrome imaging ; The Imaging Source, Germany) at 30, 300, or 600 fps and recorded directly on the HDD as uncompressed video files.

### Phototaxis Experiment

An inverted microscope (IX71) was used to control the movement direction of the cells via phototaxis. A green LED with a wavelength of 525 nm (L3-G2530-12500, FULL SUN OPTOTECH, Taiwan) served as the induction light, positioned on the microscope stage with the samples sandwiched and facing one another. To prevent *Chlamydomonas* from being attracted to the observation light owing to phototaxis, a long-pass filter (RG630, SCHOTT, Germany) with a transmission cutoff wavelength of 630 nm was placed above the condenser, and only light with wavelengths that did not induce phototaxis was used as the observation light.

### Visualization of the flow field of the liposome membrane

Color videos of chlamylipo with phase-separated membrane domains at 300 fps were split into RGB channels; the R-channel images were analyzed to track the motion of the cell body, and the G-channel images were analyzed to obtain the flow field of the liposome membrane. The centroid coordinates of the cell body were automatically measured using ImageJ software, whereas those of the phase-separated domains were determined manually. The north-south axis was defined as the swimming axis, which was calculated as the first principal component of the entire trajectory of the centroid of the cell body, and latitude and longitude were then calculated based on this axis.

### Simulation model of the membrane flow field of chlamylipos

We utilized a theory [15] to understand the relationship between the velocity field on the liposome membrane and the flagella beating force. We briefly explain here the equations used in the manuscript, as well as the assumptions underlying their derivation. Consider a spherical liposome with a membrane viscosity *η*_*m*_and radius *R*. The fluids inside/outside the membrane are incompressible Newtonian fluids with the viscosities *η*^+^ and *η*^−^, respectively. The Stokes equations that describe the velocity fields of the membrane ν^*m*^and the interior (-)/exterior (+) fluids v are given as

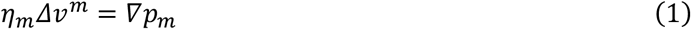

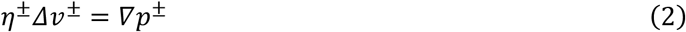

where *p*_*m*_and *p*^±^ are the pressure at the membrane and interior/exterior fluids, respectively. By assuming velocity continuity and stress balance conditions at the membrane, the velocity fields can be solved analytically when there is no deformation of the membrane. When a point force *F*_*y*_= (*0, F, 0*) is applied to the north pole (0, 0, *R*) of the spherical membrane, the velocity field at position (*R sin θ cos ϕ,R sin θ sin ϕ, R cos θ*) is given [15, 16] as :

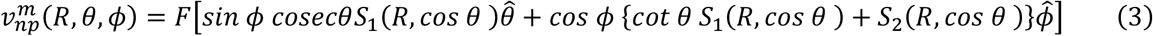

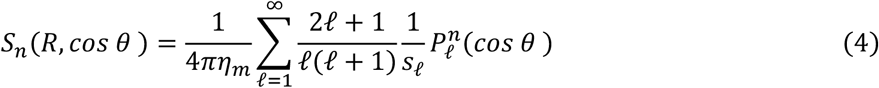

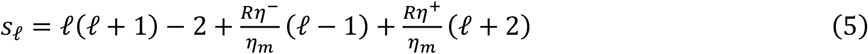

Where 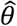 and 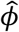 is a unit vectors in *θ* and *ϕ* directions, 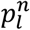 is the associated Legendre polynomial.

### Simulation model of the membrane flow field of chlamylipos

The membrane velocity resulting from the beating of two flagella can be represented by two-point forces on a spherical liposome membrane. We evaluated the exerted forces *F* and the polar angle at which the force is exerted, *θ*_*f*_, via the fitting process.

The velocity field of the membrane was evaluated by tracking the lipid domains. Since the liposome radius and the viscosities (*η*^+^ and *η*^−^) are known, we can estimate the force magnitude *F* and the polar angle *θ*_*f*_, which is the representative position where the force is exerted, from the least-squares fitting process. Because Equations (3)–(5) are the solutions under the linear Stokes equation, the velocity field owing to two-point forces can be evaluated by simply superimposing two velocity fields. By setting the force magnitude *F* and the polar angle of two forces *θ*_*f*_ as the fitting parameters, we obtain these parameters by minimizing the error of the estimated velocity and the experimentally obtained velocity. The parameters that are utilized in the fitting process are as follows: *R* = 1.0 × 10^−5^ μm, *η*^*m*^=6.5 × 10^−8^ pa . s. m [16], *η*^−^ = 1.63 × 10^−3^ pa . s, and *η*^+^ = 0.99 × 10^−3^ pa . s. The obtained force magnitude is *F* = 8.13 × 10^−11^ N and the polar angle *θ*_*f*_ = 1.41 rad from the North Pole.

## Results

### Encapsulation of *Chlamydomonas* in giant unilamellar vesicles

In the encapsulation process of *Chlamydomonas* using the liposome and water-in-oil (W/O) emulsion template method, *Chlamydomonas* was successfully encapsulated within liposomes (Fig. 1a). When observed under phase contrast, the chloroplasts of the cell body were visible inside gray liposomes (Fig. 1b). When the liposomes were observed using blue-excited green fluorescent phospholipids, both the liposome membrane and chloroplasts of the cell body were observed simultaneously (Fig. 1c). In all cases, the encapsulated cells continued to move, even when they were surrounded by liposomes. By adjusting the internal fluid with the addition of 30% Percoll to match the density of *Chlamydomonas*, the encapsulation efficiency improved, and 100.2 ± 14.5 chlamylipos/µl were achieved. In 94% of the chlamylipos, a single cell was encapsulated, and 22% of chlamilipos travelled more than twice the diameter of the liposome in 30 s while remaining enclosed in the liposome (Fig. 1d). The swimming speed of chlamylipo varied greatly, at 8.0 ± 1.9 µm/s (Fig. S1). Furthermore, no clear correlation was observed between the swimming speed and liposome diameter (Fig. S2). Despite flagellar movement, 49% of chlamylipos with a single cell rotated in place without moving forward. Chlamylipos prepared with the *Chlamydomonas pf18* mutant strain, which lacks flagellar motility, neither swimming nor rotation was observed (0.08 ± 0.05 μm/s, Movie S1). A small number of liposomes encapsulating more than two cells were observed but coordinated movement of two cells was not seen, and stable swimming was not observed (Movie S2).

**Fig. 1.**
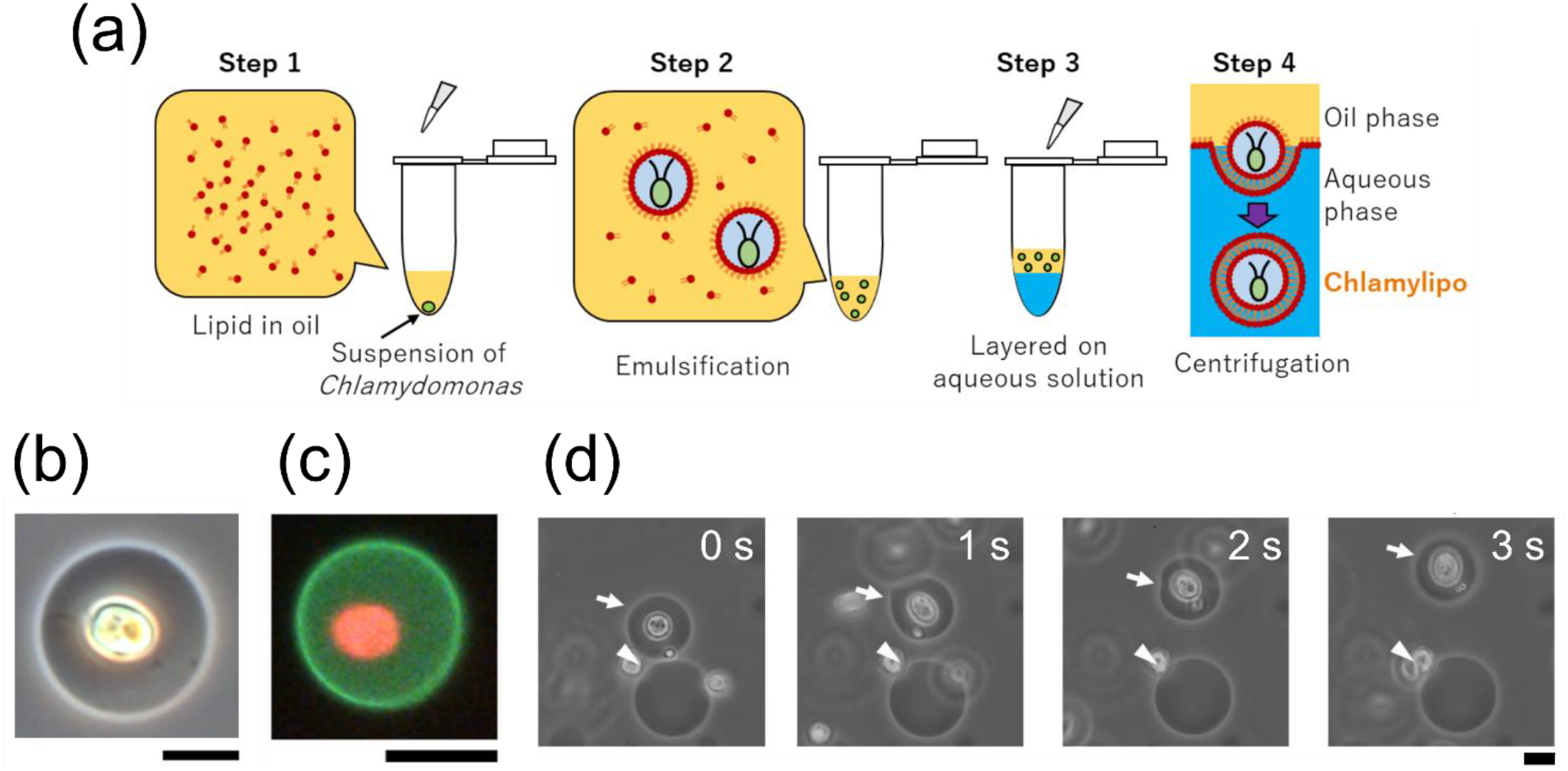
Preparation of chlamylipos, giant liposomes containing a living *Chlamydomonas*. (a) Liposomes were obtained by creating a W/O emulsion with an aqueous phase containing a phospholipid solution and *Chlamydomonas*, layered in an outer solution, and centrifuged. (b) Phase-contrast images of the produced chlamylipos (color photography). (c) Fluorescence image of a chlamylipo. Liposome membrane and chloroplast’s autofluorescence were observed green and red, respectively. (d) Time-lapse phase contrast images over three seconds of swimming chlamylipo. Liposomes without encapsulated cell did not move (indicated by an arrowhead), whereas chlamylipo moved approximately 20 μm at a constant speed in one direction over three seconds (indicated by an arrow). All scale bars represent 10 μm.

### Phototactic response of chlamylipo

When green light shone from the side during the observation of chlamylipo, it swam in the direction closer to (or further from) the light source (Fig. 2a). Subsequently, when green light shone from the opposite direction, the *Chlamydomonas* inside changed direction and swam in the opposite direction within a second (Fig. 2, Movie S3). This directional change was sustained for over 16 cycles, covering a total distance of approximately mm (Movie S4). Furthermore, even in chlamylipos that did not swim, the encapsulated *Chlamydomonas* cells repeatedly changed direction in response to green light.

**Fig. 2.**
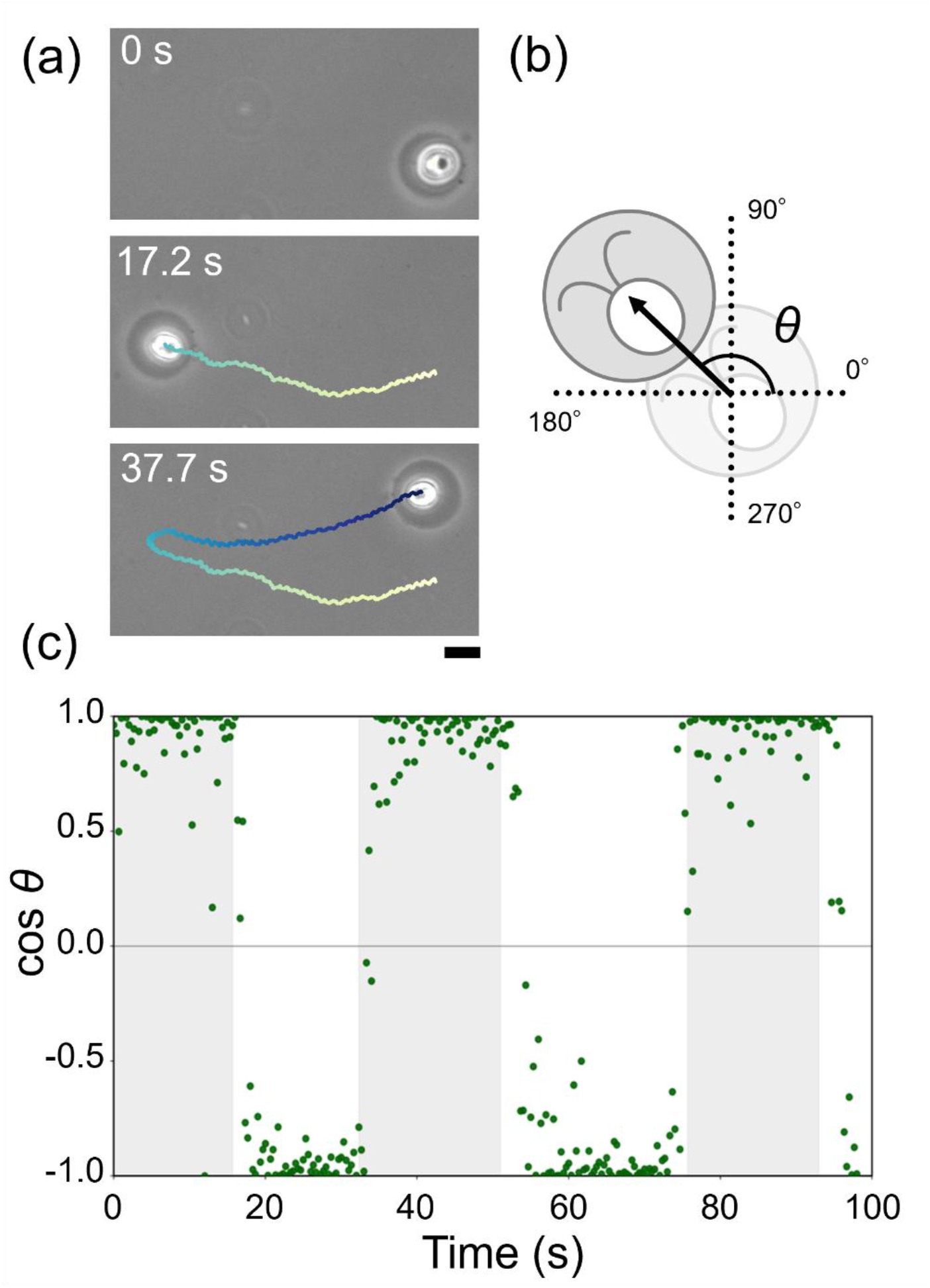
Swimming and Phototactic Direction Control of chlamylipo. (a)Phase-contrast images tracing the movement of chlamylipo. The green light was continuously illuminated from the left side of the image until 17.2 s and from the right side until 37.7 s, resulting in a change in movement direction. (b)Cosine evaluation image diagram of the chlamylipo direction. (c) Change in the direction of movement of chlamylipo. In areas with a gray background, green light was shined from the 0-degree direction (cos *θ* = 1.0), and in areas with a white background, light was shined from the 180-degree direction (cos *θ* = -1.0).

### Periodic Membrane Deformation of chlamylipo

When the liposome membrane was labeled with fluorescence and rapidly imaged using dark-field microscopy while applying excitation light, periodic membrane deformations synchronized with flagellar beating were observed (Fig. 3, Movie S5). At the beginning of the effective stroke, two membrane protrusions were formed at the front as the flagella pushed the liposome membrane outward. As the effective stroke progressed, the membrane protrusions moved from the front to the left and right. At the end of the effective stroke, as the flagellar tips were drawn close to the cell body, the membrane protrusions shortened and disappeared. During the recovery stroke, no membrane protrusions were observed, and new membrane protrusions formed at the beginning of the next effective stroke cycle. Because the plane containing the two flagella (flagellar plane) rotated around the anterior-posterior axis in conjunction with the internal rotation of *Chlamydomonas*, the length of the membrane protrusions periodically increased and decreased on the kymograph (Fig. 3b). Note that 0° in the kymograph corresponds to the direction of the *Chlamydomonas* front. In chlamylipos that did not form membrane protrusions associated with flagellar movement, forward swimming was not observed.

**Fig. 3.**
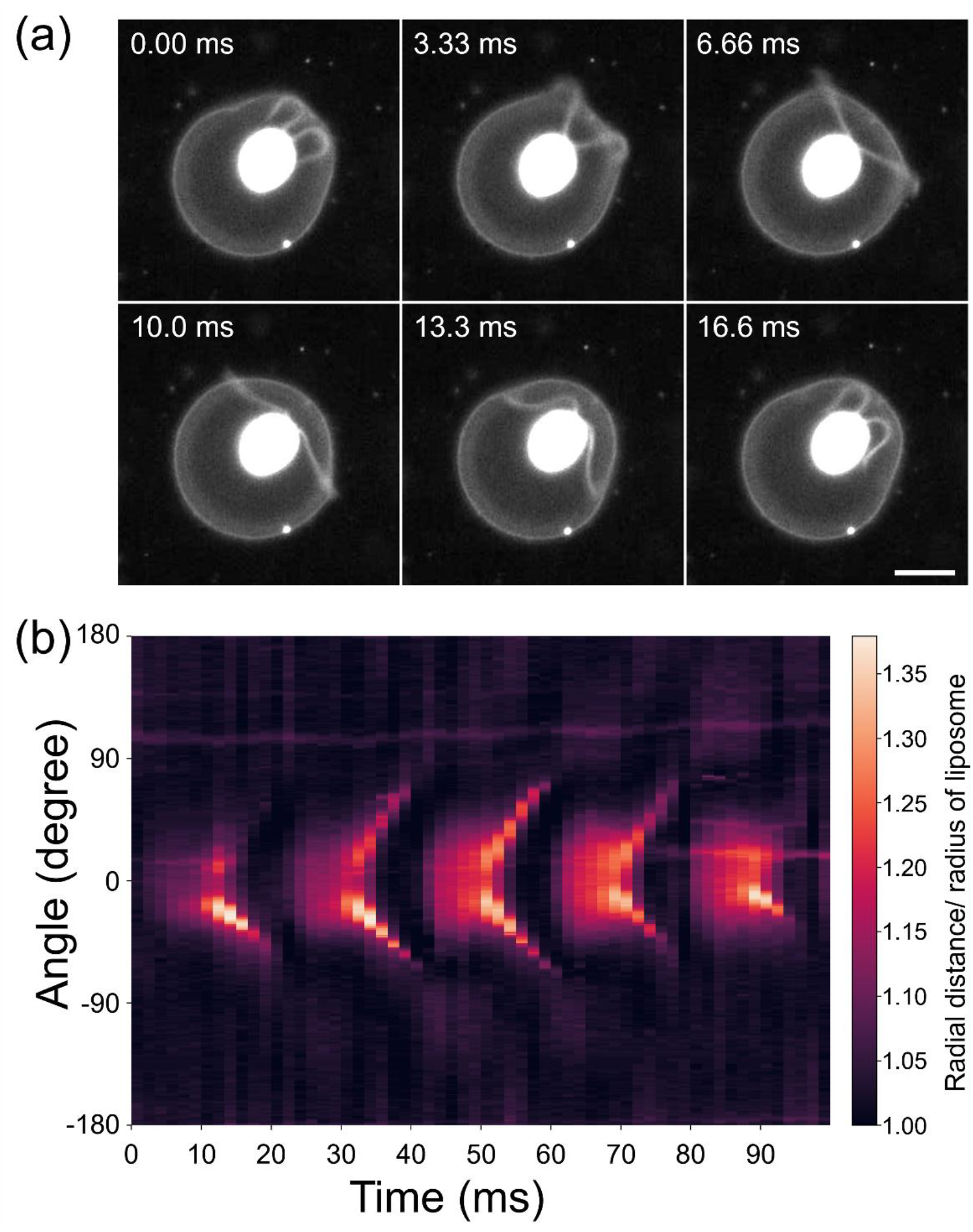
Periodic membrane deformation of a chlamylipo. (a) Superimposed image sequence of a chlamylipo recorded at 600 fps using dark-field (cell body and flagella) and fluorescence (membrane) microscopy. (b) Kymograph of the periodic membrane deformation of a chlamylipo. The radial distance of membrane elements relative to the forward direction is colour-coded.

### Coupled flow inside and outside the liposome

The flow of liquid inside and outside the chlamylipo was visualized by adding tiny fluorescent beads to either the internal or external fluid (Fig. 4). The movement of the fluorescent beads within a thickness of 1 μm, which is the focal depth of the objective lens, was observed. The tracer particles inside the chlamylipo exhibited small, rapid circular motions at several tens of hertz, while also moving back and forth along the body axis with a period of approximately 4 Hz (Movie S6-8). Similarly, the tracer particles outside the chlamylipo also showed small, rapid back-and-forth motions at several tens of hertz and moved back and forth along the body axis with a period of approximately 4 Hz (Movie 9-11).

**Fig. 4.**
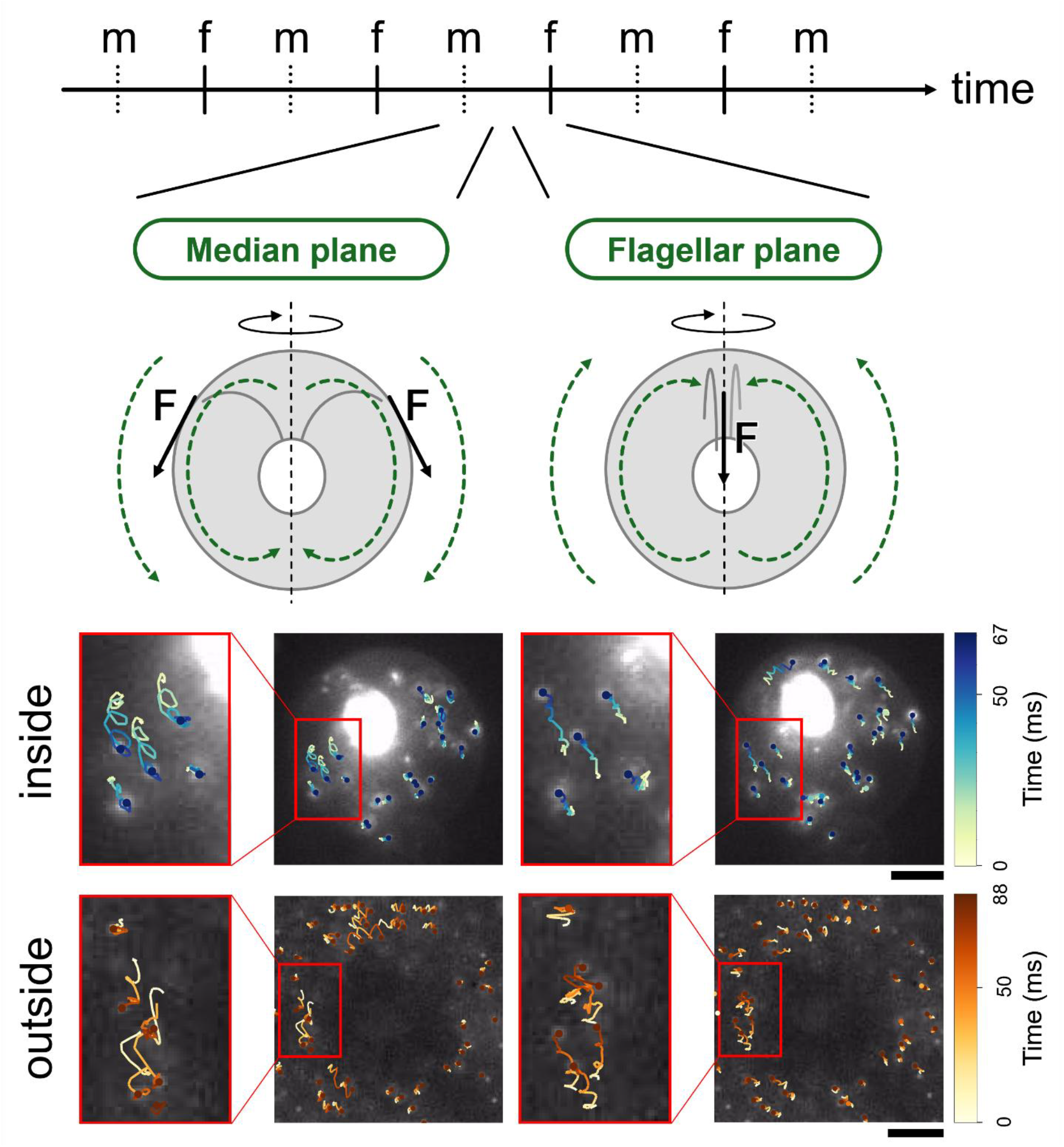
Fluid Dynamics of Internal and External Fluids in chlamylipo. As *Chlamydomonas* rotates while swimming, chlamylipo appears to alternate between two orientations when viewed perpendicular to the swimming direction: the median plane (m-plane), where the cell body is viewed from the front, and the flagellar plane (f-plane), where the flagellar beat is viewed from the side (top panels). Middle panels: Observation of tracer beads inside the liposome. The beads exhibited small, rapid circular motions at several tens of hertz. In the m-plane view, beads flow from anterior to posterior, whereas in the f-plane view, they flow from posterior to anterior. Bottom panels: Similarly, the tracer beads outside the liposome exhibited small, rapid circular motions. Beads flow from anterior to posterior in the m-plane view and from posterior to anterior in the f-plane view. The brightness of the video was adjusted to visualize the particle trajectories. Trajectories were drawn from 0 to 66.7 ms (m-plane) and 66.7 to 133.3 ms (f-plane) for the internal fluid and from 0 to 83.3 ms (m-plane) and 83.3 to 166.7 ms (f-plane) for the external fluid. Scalebar is 10 µm.

### Visualization of Membrane Dynamics Using Phase-Separated Domains

Let us consider the membrane flow generated in the liposome by the movement of *Chlamydomonas* inside a chlamylipo (Fig. 5a). The solution flow produced by the repeated effective and recovery strokes of the flagella can be approximated, as a long-term average, by two steady backward flows near the flagella. The membrane flow induced by the flagella causes the fluid inside the liposome to flow, dragging the liposome membrane along with it. Because the internal fluid is confined within the spherical liposome, the backward flagellar flow toward the rear of the cell body must always be accompanied by a return flow toward the front of the cell body. Furthermore, since the liposome membrane is a two-dimensional incompressible fluid on a closed surface, the backward membrane flow generated near the origin of the flagellar flow must also be accompanied by a return flow toward the front of the cell body. The two flagellar flows generated on the flagellar plane “f,” which contains the two flagella, are accompanied by two return flows toward the front on the median plane “m” perpendicular to it, and as a result, it is expected that four vortices will form on the liposome membrane (Fig. 5a, b). Even in the theoretical model in which point forces were applied at two points on the spherical membrane, the formation of four vortices was predicted (Fig. 5c). In reality, as *Chlamydomonas* rotates, it is thought that the four vortices also move in the equatorial direction. To visualize the membrane flow occurring on the chlamylipo, a chlamylipo was fabricated with a lipid composition that results in two-dimensional phase separation on the membrane (Fig. 5d, Movie S12). Multiple black spots representing phase-separated domains were observed on the green fluorescent liposome membrane, while the Chlamydomonas cell body rotated at 2.2 Hz. Each domain meandered or rotated on the sphere, oscillating in the anterior-posterior direction of the cell body (Fig. 5e). By subtracting the rotational speed of the cell body from the trajectories of each dot, the trajectories in a rotating coordinate system co-rotating with *Chlamydomonas* were determined. By plotting the velocity vectors of membrane flow at each point of the trajectories, a planar map of the membrane flow field as seen from *Chlamydomonas* was obtained (Fig. 5f). Rapid southward flow occurs at two locations on the sphere (Φ = -90 and 90 degrees), and at positions 90 degrees away from these, slower northward return flows are observed. Together with the east-west flows connecting the north-south flows, four vortices centered near the equator can be identified. From these results, the four-vortex structure on the liposome membrane predicted by the theoretical model was experimentally verified. Notably, domain motion was also observed in translationally swimming chlamylipos (Fig. S3,4, Movie S13), whereas it was not observed in liposomes without encapsulated cells (Fig. S5).

**Fig. 5.**
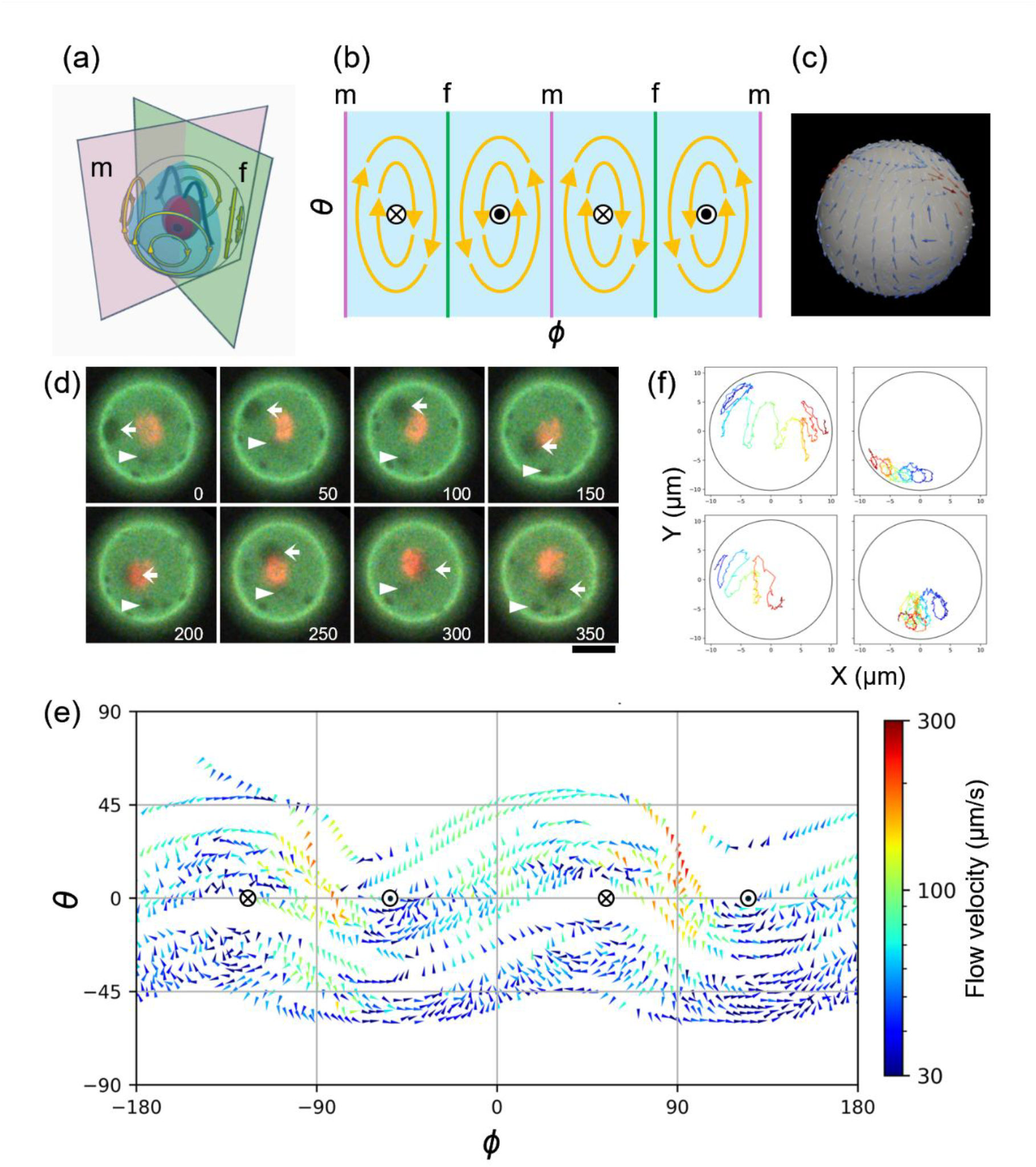
Membrane flow field on a chlamylipo. (a) Schematic diagram of a chlamylipo. Yellow arrows indicate the flow directions of four predicted vortices. Plane “f” contains two flagella of a *Chlamydomonas*, and plane “m” indicates a median plane perpendicular to the “f” plane. (b) A flat map of the membrane flow field as shown in (a). Vertical lines indicate the cutting lines on the membrane by “f” and “m” plane in (a). The centers of flor vortices are marked by ⊙ and ⊗. (c) Incompressible 2D spherical field with two-point forces. The position and orientation of the forces are consistent with *Chlamydomonas*’s flagella in (a). (d) Sequential images of a self-rotating chlamylipo. The cell body (red) and liposome membrane (green) with phase-separated domains (black dots) are shown. Elapsed times are shown in ms. Scale bar is 10 µm. (e) Typical trajectories of phase separated domains for 1 s. Time progressed from blue to red. (f) Membrane flow field in the coordinate system rotating with *Chlamydomonas* of a chlamylipo. The centers of flor vortices are marked by ⊙ and ⊗ as shown in (b).

## Discussion and Conclusion

In this study, we created “chlamylipo” by encapsulating *Chlamydomonas* inside liposomes and discovered that as the internal *Chlamydomonas* beat their flagella, the entire liposome swam forward and exhibited phototaxis. Two membrane protrusions appeared on the liposome membrane of chlamylipo in association with flagellar movement, repeating a cycle in which they moved backward and disappeared. Both the internal and external fluids of chlamylipo exhibited rapid circular or back-and-forth motions at several tens of Hz, while slowly migrating north–south at approximately 4 Hz. The membrane domains on the liposome surface also underwent small back-and-forth motions, repeatedly moving quickly southward and then slowly northward at approximately 4 Hz. From the perspective of the rotating *Chlamydomonas*, the membrane flow field was a circulating current containing four vortices, which could be reproduced using a simplified flow field model.

In the low Reynolds number environment where chlamylipo exists, microscopic objects must repeatedly undergo time-irreversible asymmetric deformations to achieve continuous center of mass movement. Chlamyipo fulfills this requirement by repeatedly forming two membrane protrusions at the front that move backward and then disappear, creating a time-irreversible asymmetric deformation cycle. It is considered that as the membrane protrusions push the surrounding water masses backward, chlamylipo as a whole moves forward in the reaction. Most chlamylipo did not swim forward, and among those that did swim, there was variation in swimming speed. No clear correlation was observed between swimming speed and liposome diameter. In addition, chlamylipo, which did not form membrane protrusions, did not move forward. Even for liposomes of the same size, variations in the length of the membrane protrusions and the distance they travel backward are presumed to affect the movement speed of chlamylipo. By examining the correlations between factors other than the liposome diameter, such as the protrusion length, duration of protrusion, and movement speed, it should be possible to clarify the factors that determine swimming speed. By doing so, we could increase both the proportion of forward-swimming chlamylipo and their swimming speed, thereby improving the efficiency of cargo transport by increasing the percentage of forward-moving chlamylipo and their forward velocity.

The internal and external liquids and membrane flow of the consisted of small back-and-forth movements at several tens of Hz and slower movements at approximately 4 Hz. Based on the frequencies of these movements, the small back-and-forth motion was attributed to flagellar movement, and the slower motion was considered to result from cell rotation. The small back-and-forth movement occurred at a frequency similar to that of flagellar motion, whereas the slower movement had a period approximately twice that of rotation. This can be explained by the fact that the flagellar surface passes over the membrane twice during one full rotation of *Chlamydomonas*. Because the internal liquid, membrane, and external liquid exhibit similar behaviors, it is believed that the internal liquid flow generated by the flagellar motion of *Chlamydomonas* is transmitted through viscous friction to the liposome membrane and external liquid, causing periodic movement.

The internal liquid of the liposome is a three-dimensional closed space with no exchange with the outside; therefore, the water mass pushed backward inevitably returns forward. Similarly, the liposome membrane is a two-dimensional closed space with no exchange with the outside; therefore, the membrane molecules pushed backward inevitably return forward. Therefore, on the flagellar plane, which includes the anterior-posterior axis of the cell body and the two flagella, a backward flow occurs, whereas on the median plane perpendicular to this, a forward return flow is thought to occur. The circulatory flow generated in the internal liquid by flagellar movement is believed to induce membrane flow through viscous friction, which, in turn, causes circulatory flow in the external liquid near the membrane.

The angular momentum generated by the oblique beating of the flagella of *Chlamydomonas* can be interpreted as being balanced by the reaction forces produced by the flows of the inner fluid, membrane, and outer fluid, allowing free-swimming *Chlamydomonas* to maintain an equivalent rotation speed. This suggests that *Chlamydomonas* inside chlamylipo exhibited phototaxis, regardless of whether forward swimming occurred.

By calculating the velocity field in the chlamylipo coordinate system, we found that *Chlamydomonas* formed a circulatory flow with four vortices. The membrane flow on the liposome surface is a two-dimensional, incompressible, viscous flow constrained by a sphere. The southward flow generated by flagellar motion must circulate and return to its original position within the two-dimensional closed region of the sphere. In chlamylipo, because the flagella contact the membrane at two points where downward flow occurs, an upward flow arises at the median plane to return to the original position, resulting in a circulatory flow with four vortices. The fastest flow velocity reached 1500 μm/s, which is comparable to the movement speed of the flagellar tip during the effective stroke.

Fitting a fluid model simulating the flow on a sphere to the measured data reproduced circulation with four vortices, capturing the velocity differences between the southward and northward currents. The total force required to generate a flow field equivalent to the measured values was 81.3 pN at both points. Previous research [17] reported a value of approximately 30 pN, suggesting that the values are of the same order of magnitude. In this study, the flow of the inner and outer fluids could only be observed in a single cross-section, and the flow in the direction perpendicular to the screen could not be detected. Because the flows of the inner fluid, membrane, and outer fluid were recorded using different chlamylipos, their temporal correlations could not be established.

Although the global membrane flow, as observed in *Chlamydomonas*, was reconstructed, the meridional and oblique flows necessary to generate rotation could not be detected in this study. Because the *Chlamydomonas* inside the chlamylipo exhibits precession motion while tilting its rotation axis, it is believed that the oblique flow cannot be accurately detected. By simultaneously observing the flows of the inner fluid, membrane, and outer fluid, as well as the morphological changes of the flagella in three dimensions within the same chlamylipo, and analyzing them after correcting for the precession motion of the cell body, it will become possible to understand the mechanism by which periodic membrane deformation determines the forward velocity and how oblique fluid motion generates rotation.

In this study, we found that chlamylipo swims forward with flagellar motion and shows phototaxis. By co-encapsulating various compounds, microparticles, or microorganisms as cargo, such as fluorescent beads used as tracers, it should be possible to transport the cargo to a target location by controlling its swimming direction with light. A more detailed understanding of membrane deformation and the flow of internal fluid, membrane, and external fluid associated with flagellar motion is expected to improve the efficiency of active drug delivery systems (DDS).

## Conflict of interest

The authors declare no conflict of interest.

## Author contributions

S.S., M.H., H.S., K.A. and T.K. designed the project. S.S., M.H., H.S. and K.A. performed the experiments and analyzed the data. S.H. and D.M conducted the fluid mechanical analysis. M.H., H.S., K.A., D.M., T.K. wrote and checked the manuscript.

## Data availability

The evidence data generated/analyzed in this study are included in this article.

## Acknowledgements

I would like to express my deepest gratitude to Professor Masafumi Hirono of the Molecular Cell Biology Laboratory at Hosei University for providing *Chlamydomonas* and technical support during this research. I would also like to extend my appreciation to Professor Taro Toyota of the University of Tokyo for his valuable suggestions regarding this study. We acknowledge the use of Gemini 3.0 pro and paperpal for enhancing the quality of the English language in this manuscript. This work was supported by JSPS KAKENHI (Grant Nos. 21H05879, 23K22673, 23H04418, and 23K26040) and JST PRESTO (Grant No. JPMJPR21OA) to D.M., and by JSPS KAKENHI (Grant Nos. 23H04430 and 21H05891) to M.H.

## Supplementary Figures

**Fig. S1.**
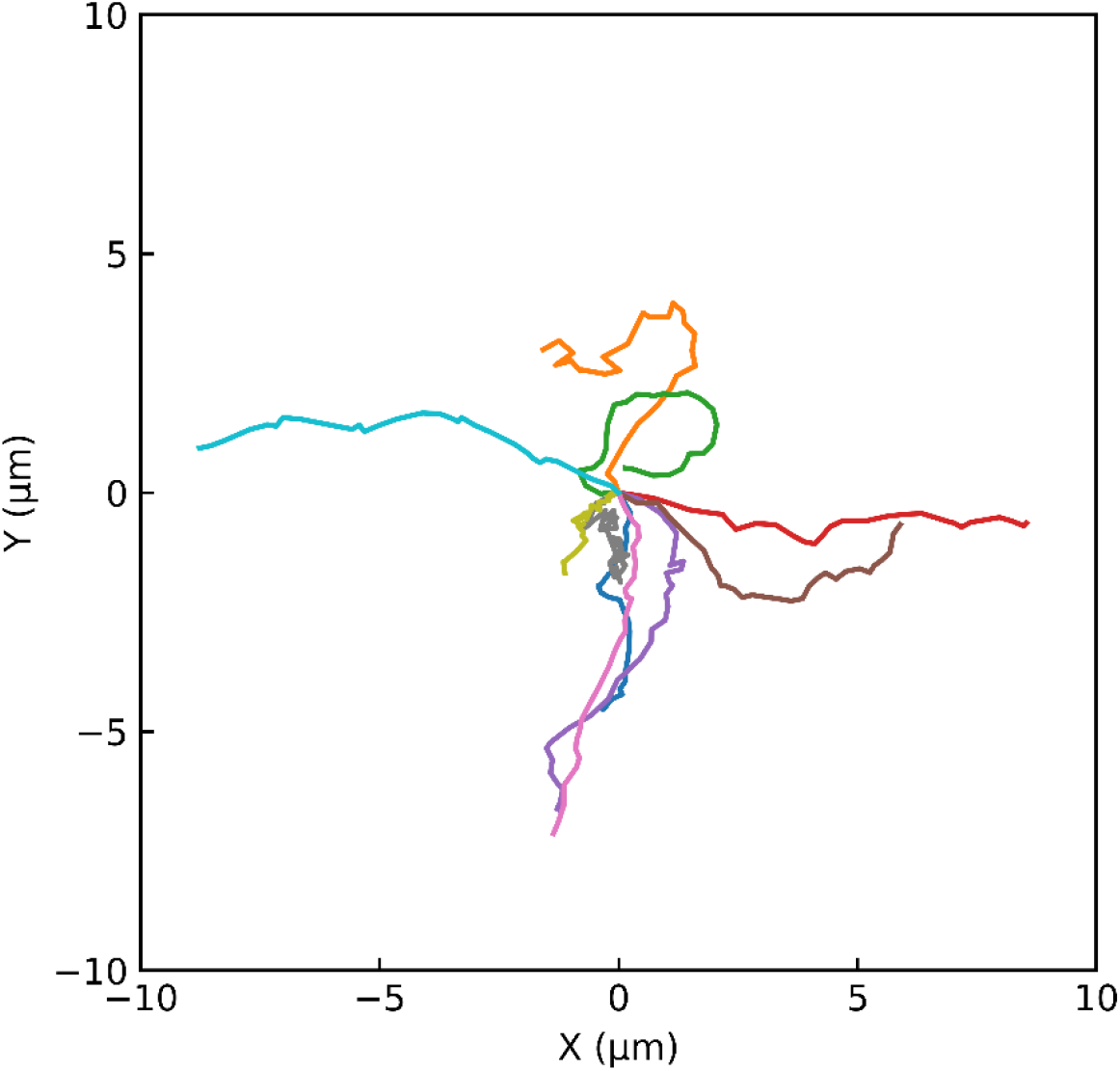
Trajectory of chlamylipos. The movement of chlamylipo was tracked for one second, starting from the origin as the initial position. The movements showed no directional patterns without photo stimulation (n=10).

**Fig. S2.**
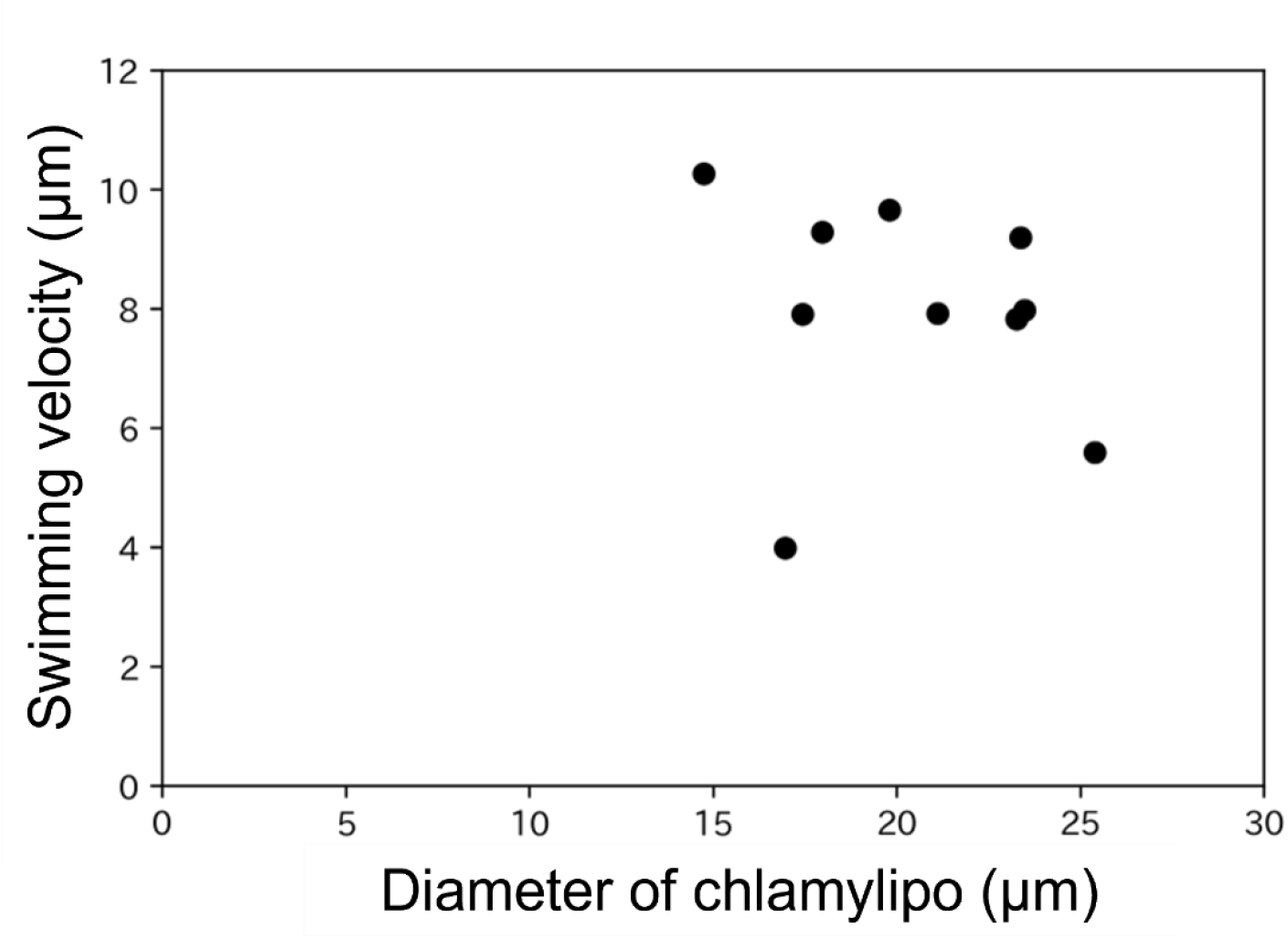
Relationship between the diameter and swimming speed of chlamylipo. The diameter and movement speed of chlamylipo are shown (n = 10). The movement speed was measured over 1 s. The diameter of the liposomes was determined as the average size of the liposomes in each frame during movement.

**Figure S3.**
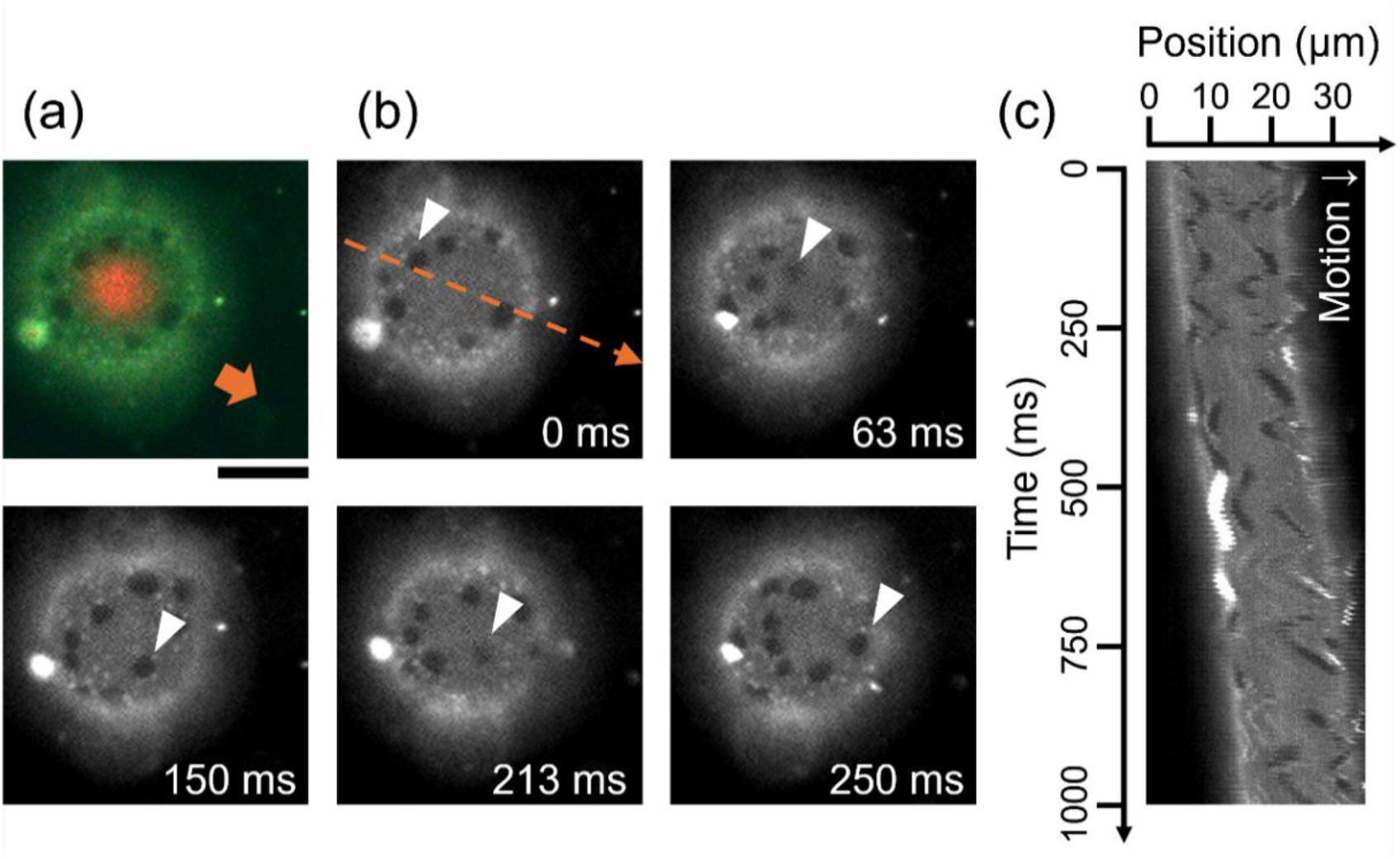
Membrane Fluidity of Swimming Phase-Separated chlamylipo. (a) Fluorescence color image of a liposome encapsulating *Chlamydomonas*. The green fluorescence represents Atto 488 DOPE in the liposome membrane, and the red fluorescence is the autofluorescence of *Chlamydomonas*. The liposome deformed and moved as the internal *Chlamydomonas* swam. (b) Time-lapse images of the green fluorescence from (a). (c) Kymograph of the dashed area in (b). The white or gray regions indicate fluorescence from the liposome membrane, while the black regions represent domains or background. The domains moved back and forth accompanying the movement of chlamylipo. Scalebar is 10 µm.

**Fig. S4.**
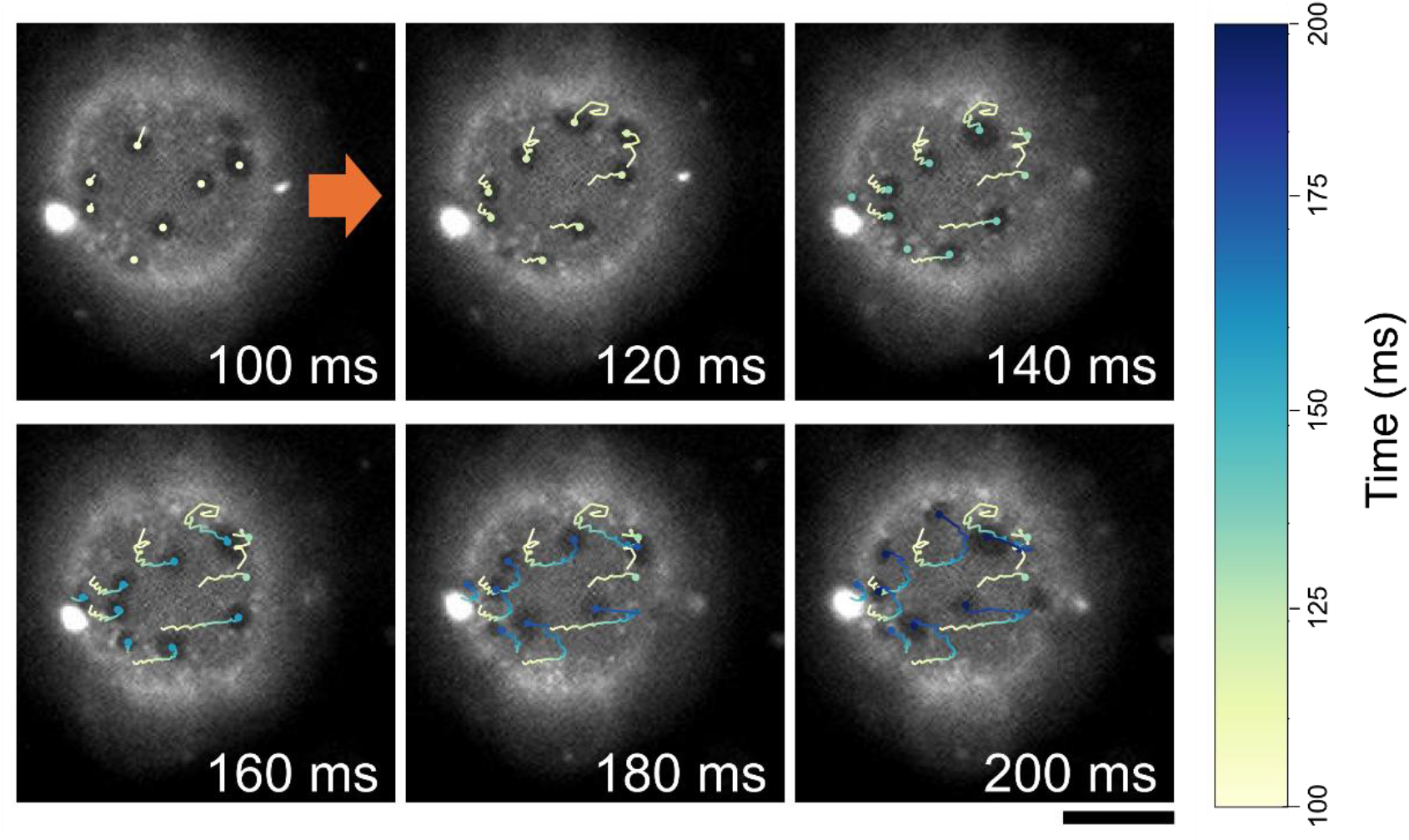
Trajectory of the domain accompanying the movement of chlamylipo. Time-lapse images of the chlamylipo domain. For chlamylipo in Fig. S3, the movement of the domain between 100 ms and 200 ms was tracked. The arrows indicate the direction of chlamylipo progression. The domain moved forward in front of the liposome and then moved backward. Scalebar is 10 µm.

**Figure S5.**
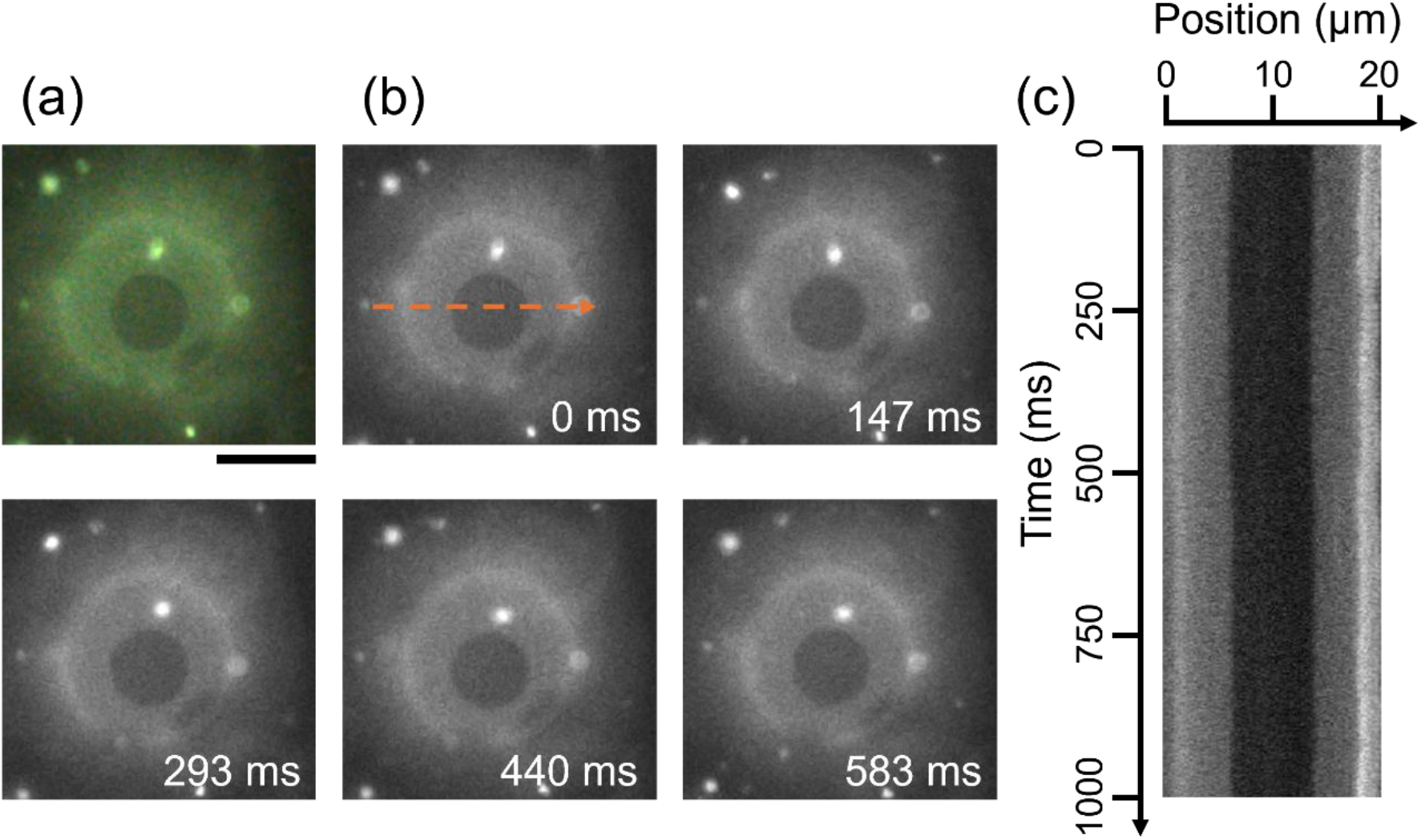
Laterally phase-separated liposomes without encapsulated *Chlamydomonas*. (a) Fluorescence color image of a phase-separated liposome. Green fluorescence corresponds to Atto 488-DOPE incorporated into the liposome. (b) Time-lapse images of the green fluorescence shown in (a). The dashed arrow indicates the line used to generate the kymograph in (c). The domains remained mostly stationary. (c) Kymograph taken along the dashed arrow in (b). White or gray regions represent the fluorescence of the liposome membrane, while black regions correspond to domains or the background.

## Movie captions

**Movie S1. Chlamylipo was prepared using non-motile *pf18* mutants**.

A liposome encapsulating the *pf18* mutant was prepared using the interface-crossing method, similar to that of wild-type chlamylipo. Because the cell lacks motility and does not induce membrane deformation, the liposome does not swim in the same manner. The video was played at real-time speed.

**Movie S2. Chlamylipo encapsulating two cells**.

Two *Chlamydomonas* cells were encapsulated within a single liposome, and their movements were not coordinated, and stable swimming was not observed. The video was played at real-time speed.

**Movie S3. Phototactic swimming control of chlamylipo**.

Video overlaid with the trajectory of the center of gravity of chlamylipo. The snapshots in Figure 2(a) were obtained from the video. In addition to the change in swimming direction in response to the switching of the green light, the trajectory exhibited a wavy pattern corresponding to the rotation of the *Chlamydomonas* cell. The video was recorded at 30 fps and played at real-time speeds.

**Movie S4. Repetitive phototactic response of chlamylipo**.

The chlamylipo shown in this video is identical to that presented in Fig. 2a and Movie S3. Chlamylipo successfully changed its swimming direction in response to repeated changes in light direction. Directional control was achieved for over 16 turns, covering a total distance of approximately 1.2 mm. The video was played at a speed of 30 ×.

**Movie S5. Periodic membrane deformation of chlamylipo**.

The chlamylipo shown in this video is identical to that shown in Fig. 3a. Chlamylipo was prepared using fluorescent phospholipids and observed under dark-field illumination while irradiated with excitation light. Periodic membrane deformations were observed using high-speed imaging at 600 fps. The video was played at 1/20× speed.

**Movie S6. Internal fluid flow in chlamylipo (m-plane)**.

The chlamylipo shown in this video is identical to that shown in Fig. S4. Fluorescent beads were co-encapsulated within chlamylipo to trace the fluid motion. High-speed imaging at 600 fps revealed periodic bead flows accompanying the flagellar beating and cell rotation. This video shows the flow in the median plane (m-plane). The video was played at 1/100× speed.

**Movie S7. Internal fluid flow in chlamylipo (f-plane)**.

This video shows the bead flow in the flagellar plane (f-plane) under the same conditions as those in Movie S6. The video was played at 1/100× speed.

**Movie S8. One-second observation of internal fluid flow**.

The video shows the internal fluid flow recorded for 1 s. The video was played at 1/20× speed.

**Movie S9. External fluid flows around chlamylipo (m-plane)**.

The chlamylipo shown in this video is identical to that presented in the bottom row of Fig. 4. Fluorescent beads were added to the external fluid to trace the motion. High-speed imaging at 600 fps revealed periodic bead flows accompanying the cell movement. This video shows the flow in the median plane (m-plane). The video was played at 1/100× speed.

**Movie S10. External fluid flows around chlamylipo (f plane)**.

This video shows the bead flow in the flagellar plane (f-plane) under the same conditions as those in Movie S9. The video was played at 1/100× speed.

**Movie S11. One-second observation of the external fluid flow**.

The video shows the external fluid flow recorded for 1 s. The video is played at 1/20× speed.

**Movie S12. Membrane fluidity of chlamylipo**.

This movie shows the same chlamylipo as shown in Fig. 5d. Lateral phase separation occurs in the liposomal membrane, allowing the observation of membrane fluidity via domain dynamics. High-speed imaging (300 fps) revealed domain displacement induced by cell motility. Green indicates membrane fluorescence and red indicates chloroplast autofluorescence in *Chlamydomonas*. The video was played at 1/20× speed.

**Movie S13. Membrane fluidity of swimming chlamylipo**.

This movie shows the observation of membrane fluidity in swimming chlamylipo, which exhibits lateral phase separation in its liposomal membrane upon exposure to a hypertonic solution. Similar to chlamylipo (Fig. 5), back-and-forth motion of the domains accompanying flagellar beating was observed. Green indicates membrane fluorescence and red indicates chloroplast autofluorescence in *Chlamydomonas*. The video was played at 1/20x speed.

## Notes

### Competing Interest Statement

The authors have declared no competing interest.

